# Engineering of membrane complex sphingolipids improves osmotic tolerance of *Saccharomyces cerevisiae*

**DOI:** 10.1101/780817

**Authors:** Guoxing Zhu, Nannan Yin, Qiuling Luo, Jia Liu, Xiulai Chen, Liming Liu, Jianrong Wu

## Abstract

In order to enhance the growth performance of *S. cerevisiae* under harsh environmental conditions, mutant XCG001, which tolerates up to 1.5M NaCl, was isolated via adaptive laboratory evolution (ALE). Comparisons made via transcriptome data of XCG001 and the wild-type strain identified *ELO2* as being associated with osmotic tolerance. Overexpression of *ELO2* increased the contents of inositol phosphorylceramide (IPC, t18:0/26:0), mannosylinositol phosphorylceramide (MIPC, t18:0/22:0(2OH)), MIPC (d18:0/22:0), MIPC (d20:0/24:0), mannosyldiinositol phosphorylceramide (M(IP)_2_C, d20:0/26:0), M(IP)_2_C (t18:0/26:0(2OH)) and M(IP)_2_C (d20:0/26:0(2OH)) by 88.3-, 166.9-, 63.3-, 23.9-, 27.9-, 113.8- and 208.1-fold at 1.0 M NaCl, respectively, compared those of strain XCG002. As a result, membrane integrity, cell growth and cell survival of the *ELO2* overexpression strain (XCG010) increased by 24.4%, 29% and 22.1% at 1.0 M NaCl, respectively, compared those of strain XCG002. The findings provided a novel strategy for engineering complex sphingolipids to enhance osmotic tolerance.

**IMPORTANCE:** This study demonstrated a novel strategy for manipulation membrane complex sphingolipids to enhance *S. cerevisiae* tolerance to osmotic stress. Osmotic tolerance was related to sphingolipid acyl chain elongase, Elo2, via transcriptome analysis of the wild-type strain and an osmotic tolerant strain generated from ALE. Overexpression of *ELO2* increased complex sphingolipid with longer acyl chain, thus improved membrane integrity and osmotic tolerance.

## INTRODUCTION

During industrial fermentation, growth performance (concentration and growth rate) of industrial strains is a key factor affecting the efficiency of the fermentation process (1). Growth performance is determined by physiological characterization as well as nutritional and environmental conditions. Concentration and growth rate declined when industrial strains were subjected to harsh environmental conditions (2, 3). The membrane is a natural barrier separating extracellular conditions from intracellular components (4). Therefore, improving membrane function is a potential strategy to enhance the growth performance of industrial strains under harsh industrial conditions (5).

Microbial membranes are primarily composed of a mix of proteins and lipids, and manipulating these components may regulate the physiological function and growth of industrial strains (5). Cell membrane proteins have a number of different functions, but membrane protein engineering is focused on transporters as follows (3): (i) Engineering influx transporters: Expressing influx transporters is a strategy used to increase carbon flux for product synthesis (6). For example, when GatA, a transporter protein from *Aspergillus niger*, was expressed in *S. cerevisiae*, D-galUA was used as a substrate, and the meso-galactaric titer was increased to 8.0 ± 0.6 g L^−1^ (7); (ii) Engineering efflux transporters: Efflux transporters move intracellular molecules across the cell membrane to maintain cell homeostasis and eliminate waste. For instance, when human fatty alcohol transporter Fatp1 was expressed in yeast, the secretion efficiency of fatty alcohol was improved by 5-fold, and the yeast tolerance to fatty alcohol was also increased (8); (iii) Directed evolution of transporters: The efficiency and function of the original transporter protein is usually insufficient to meet the demands of industrial production. Therefore, properties and expression levels of targeted transporters may be altered via protein evolution strategy (6, 9), an example being the nicotinamide riboside membrane transporter, PnuC in *E. coli*, which was successfully engineered to accept thiamine as a substrate whereby thiamine transport efficiency was improved to 2.5 μM by directed evolution(10).

Manipulation of membrane lipids, such as phospholipids, sphingolipids and sterols, is also an efficient strategy to enhance membrane functions (11). Based on the structure of phospholipids, one engineering strategy is to modulate phospholipid head groups by altering the expression of key phospholipid biosynthesis enzymes (12). A another strategy is to regulate phospholipid fatty acid tail by changing fatty acid length, via increasing the ratio of saturated to unsaturated fatty acid and producing trans unsaturated fatty acids (tufa) (13-15). For example, through expression of cis-trans isomerase (Cti) from *Pseudomonas aeruginosa*, the tufa were incorporated into the *E. coli* membrane, decreasing membrane fluidity, as a result of which robustness and the bio-renewable fuel titer were improved (16). The content and compositions of sterols can be changed by engineering key enzymes associated with sterol biosynthesis or by changing the transcription levels of sterol biosynthesis enzymes, which are affected by global transcription factors, such as Upc2 and Ecm22, and mediators, such as CgMed3AB (17-19). For example, the expression of a key sterol C-5 desaturase FvC5SD from an edible mushroom in fission yeast, improved the ergosterol and oleic acid contents, which resulted in enhanced tolerance to ethanol and high temperature (20). Sphingolipid metabolism is regulated by sphingolipid biosynthesis key enzymes and the target of Rapamycin Complex 1 (TORC1) signals (11, 21). Thus, sphingolipid metabolism can be manipulated by TORC1, as well as the sphingosine backbone and acyl chain (22). Some attempts had been made to change sphingolipid content via engineering or the simulation of molecular dynamics (23-25). An increase in sphingolipids resulted in a *Zygosaccharomyces bailii* membrane, which was more packed and dense and increased acetic acid resistance (24). These findings indicated that the importance of developing novel strategies to improve stress resistance by engineering complex sphingolipid metabolism.

In this study, the length of sphingolipid acyl chain was engineered to change complex sphingolipid metabolism and to increase the osmotic tolerance of *S. cerevisiae*. Adaptive laboratory evolution and RNA-seq analysis led to the selection of *ELO2*, a sphingolipid acyl chain elongase. Overexpression of *ELO2* changed fatty acid, phospholipid, and complex sphingolipid contents, resulting in improved cell membrane integrity.

## RESULT

### Globe-transcriptome analysis of the adaptive laboratory evolution strain and wild-type strain

In order to understand how *S. cerevisiae* adapts to higher osmotic stress, ALE was utilized to generate osmotic tolerance mutants. The concentration of NaCl was increased with time in a stepwise fashion, to reach 1.5 M (Fig. 1A). After 300 generations of ALE, a clone (mutant XCG001) was isolated from the evolved population. The osmotic sensitivity of mutant XCG001 and the wild-type strain were tested, where the IC_50_ value of the wild-type strain and mutant XCG001 were 0.94 M and 1.40 M NaCl, respectively (Fig. 1B). The final biomass of mutant XCG001 was similar to that of the wild-type strain at 0 M NaCl, whereas the final biomass of mutant XCG001 increased by 346.5% compared with that of the wild-type strain at 1.5M NaCl (Fig. 1C and D).

**FIG. 1.**
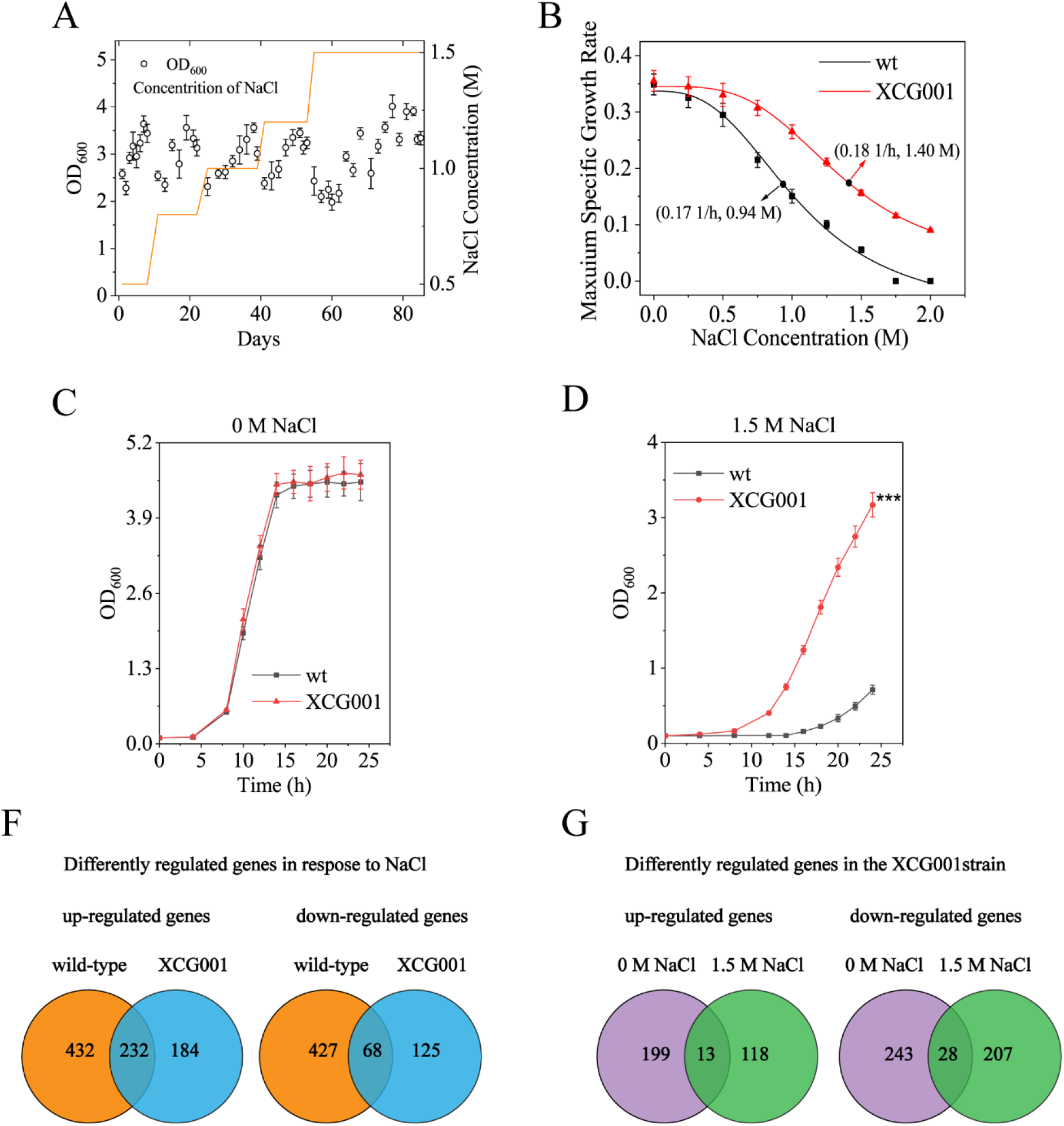
Globe-transcriptome analysis of the mutant XCG001 and the wild-type strain. (A) Cell growth trajectory showing changes in fitness during the ALE in YNB medium with different concentration of NaCl. Concentration of NaCl was stepwise improved from 0.5 to 1.5 M over time (orange line). (B) Maximum specific exponential growth rates of wild-type strain and mutant XCG001 in YNB medium supplemented with increasing concentrations of salt. The IC_50_ was calculated by the fitting curve to the data. (C) Growth profiles of mutant XCG001 and the wild-type strain in minimal medium under 0 M NaCl condition. (D) Growth profiles of mutant XCG001 and the wild-type strain in YNB medium under 1.5 M NaCl condition. (F) Venn diagrams depicting the numbers of upregulated and downregulated genes in the wild-type strain and mutant XCG001 under 1.5 M NaCl condition compared with those genes expression levels in the corresponding strains under the 0 M NaCl condition. (G) Numbers of upregulated and downregulated genes in mutant XCG001 relative to their expression in the wild-type strain under 0 M and 1.5 M NaCl conditions.

To identify the differentially regulated genes contributing to osmotic tolerance in mutant XCG001, the transcriptome sequencing (RNA-seq) was conducted to compare global gene expression in mutant XCG001 and the wild-type strain at 0 M and 1.5 M NaCl. Restrictive thresholds [|log_2_(fold change)| ≥1.5; false-discovery rate (FDR), <0.05] of significantly expressed genes were used to screen these genes. First, the differentially expressed genes were analyzed, at 1.5 M NaCl relative to 0 M NaCl in both the wild-type strain and mutant XCG001 (Fig. 1F). Transcriptional profiling analysis revealed that the expression levels of 1159 genes were significantly changed in the wild-type strain, where 664 were up-regulated and 495 were down-regulated, while in mutant XCG001, the expression levels of 609 genes displayed differential expression, where 416 were up-regulated and 193 were down-regulated. Additionally, 229 up-regulated and 68 down-regulated genes were common to both strains. Gene Ontology (GO) analysis indicated that the commonly up-regulated genes were involved in glycolysis/gluconeogenesis, pyruvate metabolism, lipid metabolism, signaling transduction, fructose and mannose metabolism. On the other hand, the 68 down-regulated genes were involved in ribosome and amino acid metabolism (supplemental Data Sets S1 and S2).

Next, significantly expressed genes in mutant XCG001, relative to those in the wild-type strain, were analyzed at both 0 M and 1.5 M NaCl (Fig. 1G). At 0 M NaCl, the expression levels of 212 genes were up-regulated and 271 genes were down-regulated. At 1.5 M NaCl, 131 genes were up-regulated and 235 genes were down-regulated. Based on GO analysis, 13 genes that were commonly up-regulated at 0 M and 1.5 M NaCl were involved in transport, pyrimidine metabolism and lipid metabolism, whereas 28 commonly down-regulated genes were involved in pyruvate metabolism and transport (supplemental Data Sets S3 and S4). These results suggested that mutant XCG001 strengthened transport, pyrimidine metabolism and lipid metabolism, which contribute to osmotic tolerance.

### Overexpression of *ELO2* enhanced osmotic tolerance

Above results showing that 13 genes were commonly up-regulated in the mutant XCG001 relative to those of the wild-type strain at both 0 M and 1.5 M NaCl, indicated that these genes may contribute to osmotic tolerance. Initially, the mRNA levels of the 13 genes were further verified by quantitative reverse transcription-PCR (qRT-PCR) analysis (Fig. S1). Next, these genes were overexpressed in each strain and evaluated for resistance to osmotic stress (Fig. S2). Interestingly, only overexpression of *ELO2* conferred resistance to osmotic stress. To confirm whether expression of *ELO2* was positively correlated with osmotic tolerance, *ELO2* was overexpressed with two other constitutive promoters, P*_TDH3_* (promoter activity weaker than that of P*_TEF1_* (26)) and P*_ADH1_* (promoter activity weaker than that of P*_TDH3_*). Although the spot results showed no obvious difference among the strains P*_ADH1_–ELO2* (XCG016), P*_TDH3_–ELO2* (XCG017) and P*_TEF1_–ELO2* (XCG010) at 1.0 M NaCl (Fig. 2A), the IC_50_ value of the strains XCG016, XCG017 and XCG010 were 1.15 M, 1.18 M and 1.22 M, respectively (Fig. 2B). The growth curves of those four strains were also distinct (Fig. 2C and D). At 0 M NaCl, the final biomass of strains XCG016, XCG017 and XCG010 was similar to that of strain XCG002 (wild-type strain containing a control plasmid pY13), whereas at 1.0 M NaCl, the final biomass of strains XCG016, XCG017 and XCG010 increased by 23%, 26% and 29%, respectively, compared to that of strain XCG002 (Fig. 2C and 2D). In addition, survival curves were generated for the four strains over a broad concentration range of NaCl (Fig. 2E). At 1.0 M NaCl, 59.5% of strain XCG002 survived, while strains XCG016, XCG017 and XCG010 exhibited 70.2%, 71.9% and 72.6% survival, indicating approximate increases of 18.0%, 20.8% and 22.1%, respectively. These results suggested that the overexpression of *ELO2* enhanced osmotic tolerance of *S. cerevisiae*.

**FIG. 2.**
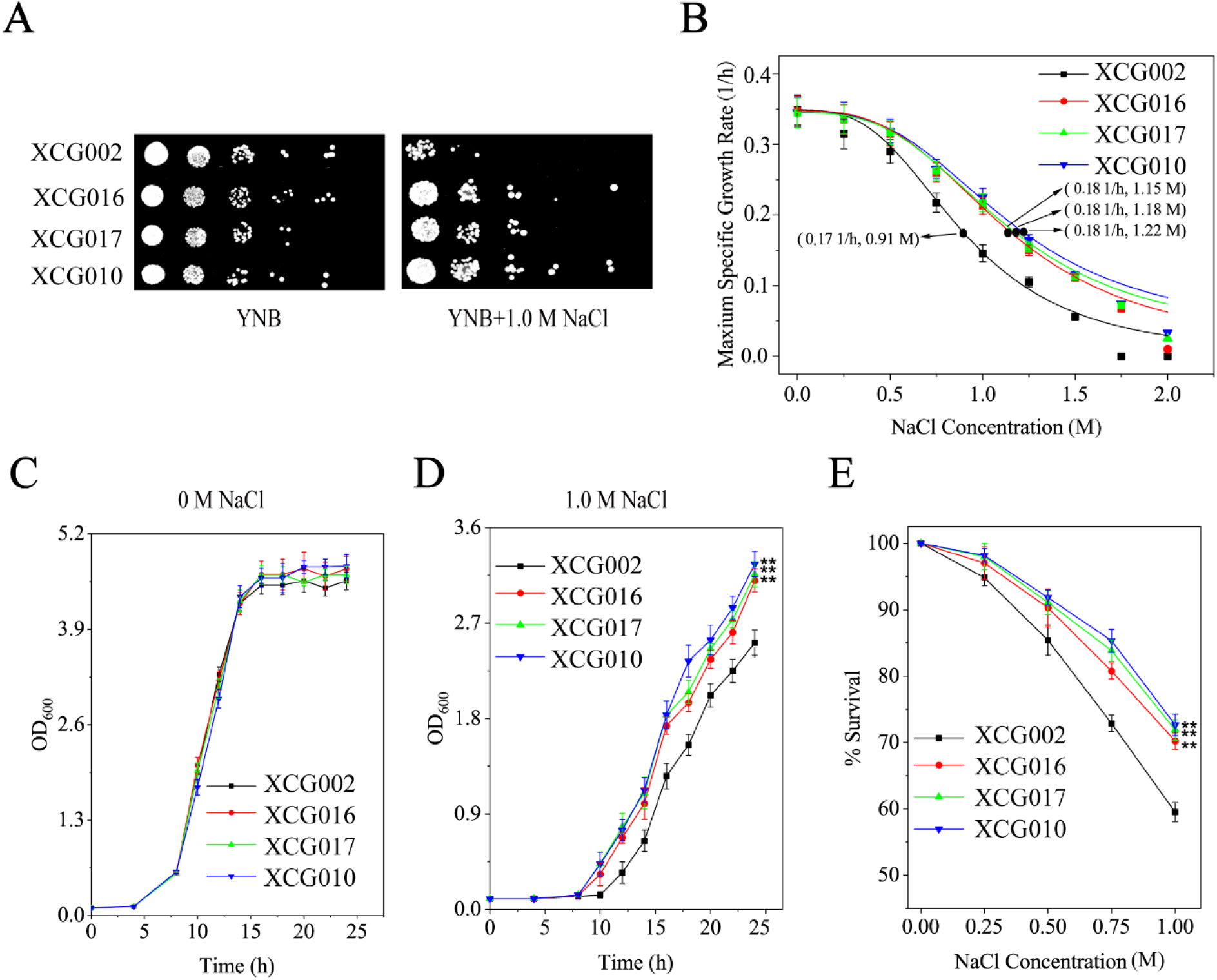
Overexpression of *ELO2* enhanced osmotic tolerance. (A) Strains XCG002, XCG016, XCG017 and XCG010 were spotted on YNB plates at 0 M and 1.0 M NaCl. (B) Maximum specific exponential growth rates of strains XCG002, XCG016, XCG017 and XCG010 in YNB medium supplemented with increasing NaCl concentration. The IC_50_ was calculated by the fitting curve to the data. (C and D) Growth curves of strains XCG002, XCG016, XCG017 and XCG010 at 0 M and 1.0 M NaCl. (E) The survival rates of strains XCG002, XCG016, XCG017 and XCG010 over of a range of NaCl doses (0.00, 0.25, 0.50, 0.75, 1.00 M). All data are presented as mean values of three independent experiments. Error bars indicate the standard deviations. *, *P* < 0.05; **, *P* <0.01; ***, *P* <0.001.

### Overexpression of *ELO2* enhanced very long fatty acids contents

The fatty acids contents of the strains XCG016, XCG017, XCG010 and XCG002 were analyzed by gas chromatography. Altered membrane fatty acids, especially C22:0, in strains XCG016, XCG017, and XCG010, compared to those of strain XCG002 at 0 M or 1.0 M NaCl are shown (Fig. 3A and B). At 0 M NaCl, the contents of C20:0, C22:0 and C24:0 in strain XCG016 increased by 32.5%, 70.8% and 12.7% compared to those in strain XCG002, respectively, whereas the level of C16:0, C16:1, C18:0 and C18:1 remained unchanged. In strain XCG017, the contents of C20:0, C22:0 and C24:0 increased by 40.9%, 78.1% and 19.0%, respectively, while the contents of C16:0, C16:1, C18:0 and C18:1 remained unchanged. In strain XCG010, the contents of C20:0, C22:0 and C24:0 increased by 52.3%, 94.1% and 14.4%, respectively, whereas the contents of C16:0, C16:1, C18:0 and C18:1 remained the same. At 1.0 M NaCl condition, the contents of C20:0 and C22:0 in strain XCG016 increased by 21.5% and 90.3% compared to those of strain XCG002, respectively, whereas the level of C16:0, C16:1, C18:0 and C18:1 remained unchanged. In strain XCG017, the content of C20:0, C22:0 and C24:0 increased by 20.3%, 26.0% and 95.8%, respectively, whereas the contents of C16:1, C18:0, C18:1 and C16:0 remained unchanged. In strain XCG010, the contents of C20:0, C22:0 and C24:0 increased by 33.1%, 106.4% and 31.5%, respectively, while the contents of C16:0, C16:1, C18:0 and C18:1 remained unchanged. Those results indicated the change in fatty acid content in XCG016, XCG017, and XCG010 was similar at 0 M and 1.0 M NaCl.

**FIG. 3.**
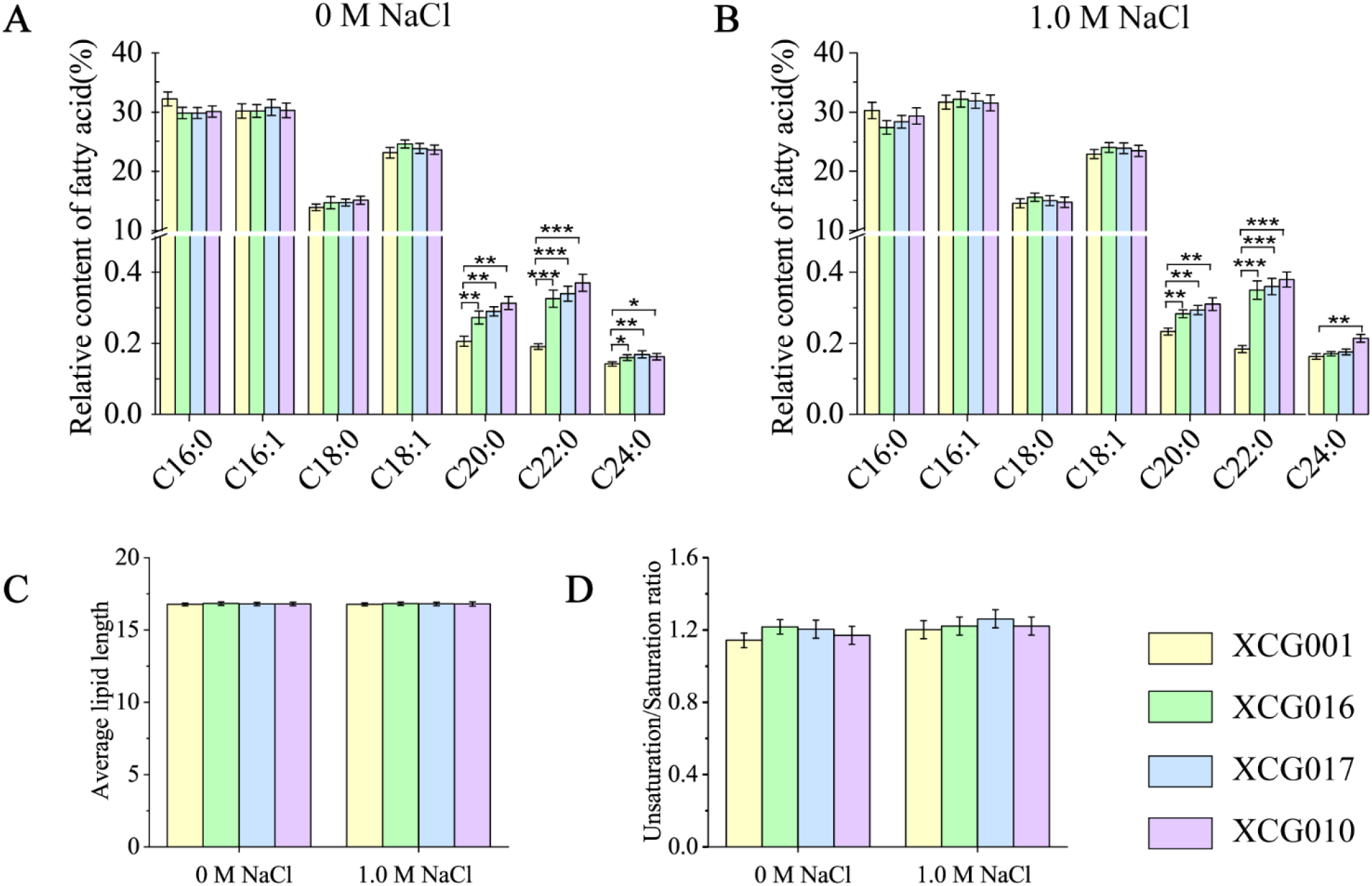
Overexpression of *ELO2* enhanced very long fatty acids contents. (A) Fatty acids contents changes in strains XCG002, XCG016, XCG017 and XCG010 at 0 M NaCl. (B) Fatty acids contents changes in strains XCG002, XCG016, XCG017 and XCG010 at 1.0 M NaCl. (C) Fatty acid average length of strains XCG002, XCG016, XCG017 and XCG010 at 0 M and 1.0 M NaCl. (D) U/S ratio of strains XCG002, XCG016, XCG017 and XCG010 at 0 M and 1.0 M NaCl. All data are presented as mean values of three independent experiments. Error bars indicate the standard deviations. *, *P* <0.05; **, *P* <0.01; ***, *P* <0.001.

The average fatty acid length in strains XCG016, XCG017 and XCG010 was almost identical to that of strain XCG002 at 0 M or 1.0 M NaCl, suggesting that the membrane “thickness” was not affected by overexpression of *ELO2* (Fig. 3C). The reason for the lack of change in membrane “thickness” may be that, although C20:0 and C22:0 contents increased by approximately 50%-100%, the proportion of C20:0 and C22:0 content to the total fatty acid content was only approximately 0.5%. In addition, the fatty acid unsaturation/saturation (U/S) ratio did not increase in the strains XCG016, XCG017 or XCG010 at 0 M or 1.0 M NaCl (Fig. 3D).

### Overexpression of *ELO2* altered complex sphingolipids contents

To test whether overexpression of *ELO2* altered the levels of phospholipids and complex sphingolipids, strain XCG010 (the strain which displayed the best osmotic tolerance) was selected to further analyze membrane phospholipids and complex sphingolipids. Overexpression of *ELO2* changed the phospholipid contents in strain XCG010 (Fig. 4A-F). At 0 M NaCl, phosphatidylserine (PS) content in strain XCG010 increased by 11.0% compared to that of strain XCG002, whereas phosphatidic acid (PA) contents decreased by 15.2%, and phosphatidylinositol (PI), phosphocholine (PC), phosphatidylglycerol (PG) and phosphatidylethanolamine (PE) contents remained unchanged. At 1.0 M NaCl, PE and PS contents in strain XCG010 increased by 15.0% and 18.9%, respectively, while PC and PA contents remained unchanged, and PI and PG contents decreased by 10.2% and 40% compared to those of strain XCG002, respectively. Further, overexpression of *ELO2* changed the complex sphingolipids contents in strain XCG010 (Fig. 4G). At 0 M NaCl, the contents of IPC (t18:0/26:0), MIPC (t18:0/22:0(2OH)), MIPC (t20:0/22:0(2OH)), MIPC (d18:0/22:0) and MIPC (d18:0/26:0) in strain XCG010 increased by 48.7-, 45.5-, 31.1-, 55.0- and 40.8-fold, respectively, but the contents of MIPC (t20:0/26:0), MIPC (t18:0/20:0(2OH)), MIPC (d18:0/24:0) and M(IP)_2_C (d20:0/26:0(2OH)) in strain XCG010 decreased by 96.5%, 99.1%, 97.3% and 99.7%, respectively, compared with the corresponding value in strain XCG002 (Fig. 4G). At 1.0 M NaCl, the contents of IPC (t18:0/26:0), MIPC (t18:0/22:0(2OH)), MIPC (d18:0/22:0), MIPC (d20:0/24:0), M(IP)_2_C (d20:0/26:0), M(IP)_2_C (t18:0/26:0(2OH)) and M(IP)_2_C (d20:0/26:0(2OH)) in strain XCG010 increased by 88.3-, 166.9-, 63.3-, 23.9-, 27.9-, 113.8- and 208.1-fold, respectively, whereas the contents of IPC (d18:1/22:0), MIPC (t16:0/18:0), MIPC (t16:0/18:0(2OH)), MIPC (t16:0/20:0(2OH)), MIPC (t18:0/20:0(2OH)) and MIPC (d20:0/26:0) in strain XCG010 decreased by 96.7%, 96.3%, 99.7%, 95.7%, 99.3% and 88.0%, respectively, compared with the corresponding value in strain XCG002 (Fig. 4G).These results suggest that high levels of complex sphingolipids with the longer acyl chain maybe enhance the osmotic tolerance.

**FIG. 4.**
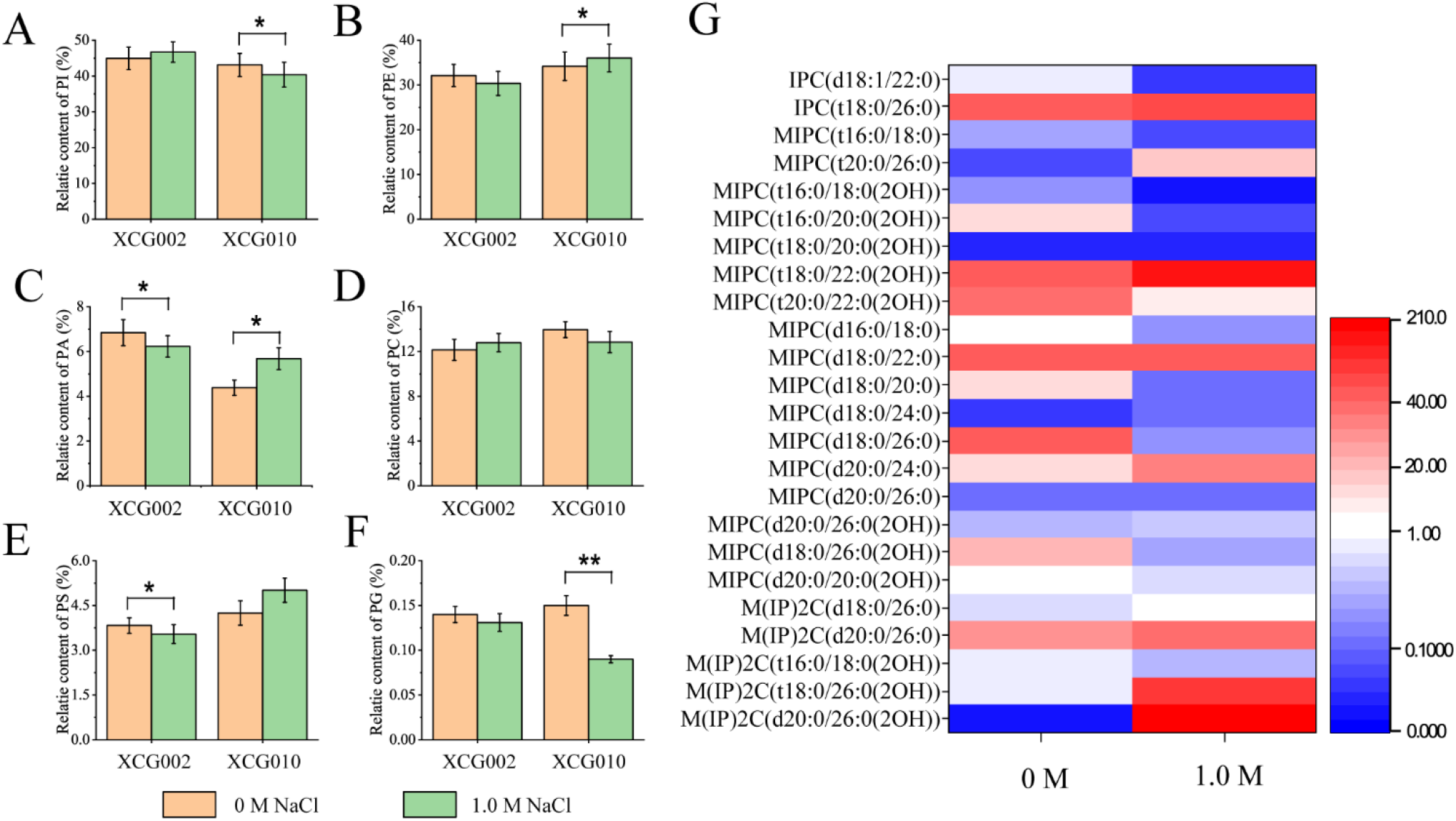
Overexpression of *ELO2* changed complex sphingolipids contents. (A-F) Phospholipid content (including PI, PE, PA, PC, PS and PG) changed in strain XCG002 and strain XCG010 at 0 M and 1.0 M NaCl. (G) Ratio of complex sphingolipid content in strain XCG010 to that of strain XCG002 changed at 0 M and 1.0 M NaCl. All data are presented as mean values of three independent experiments. Error bars indicate the standard deviations. *, *P* <0.05; **, *P* <0.01.

### Complex sphingolipids improve osmotic tolerance

To validate whether an increase of complex sphingolipids contents enhanced osmotic tolerance, the genetic details of strain XCG002 and XCG010 were investigated. The mRNA expression levels of the complex sphingolipid biosynthesis genes in strain XCG002 and XCG010 were compared at 0 M and 1.0 M NaCl. At 0 M NaCl, the mRNA levels of *AUR1, CSG2, IPT1, LAG1* and *LAC1* in strain XCG010 were increased by 1.4-, 1.7-, 1.3-, 1.5- and 1.8-fold, compared to the corresponding value of strain XCG002 (Fig. 5B). At 1.0 M NaCl, mRNA levels of *AUR1, CSG2, IPT1, LAG1* and *LAC1* in strain XCG010 were increased by 1.5-, 2.8-, 1.5- 2.1- and 2.5-fold, compared to the corresponding value of strain XCG002 (Fig. 5C). These results are consistent with the high content of complex sphingolipids detected in strain XCG010 under osmotic stress. However, the mRNA levels of the complex sphingolipid biosynthesis genes was different from the comparison between mutant XCG001 and the wild-type strain (Table S1). This was because the objects (strain XCG010 to XCG002; mutant XCG001 to the wild-type strain) and conditions (under 1.0 M NaCl and 1.5 M NaCl) of comparison were different.

**FIG. 5.**
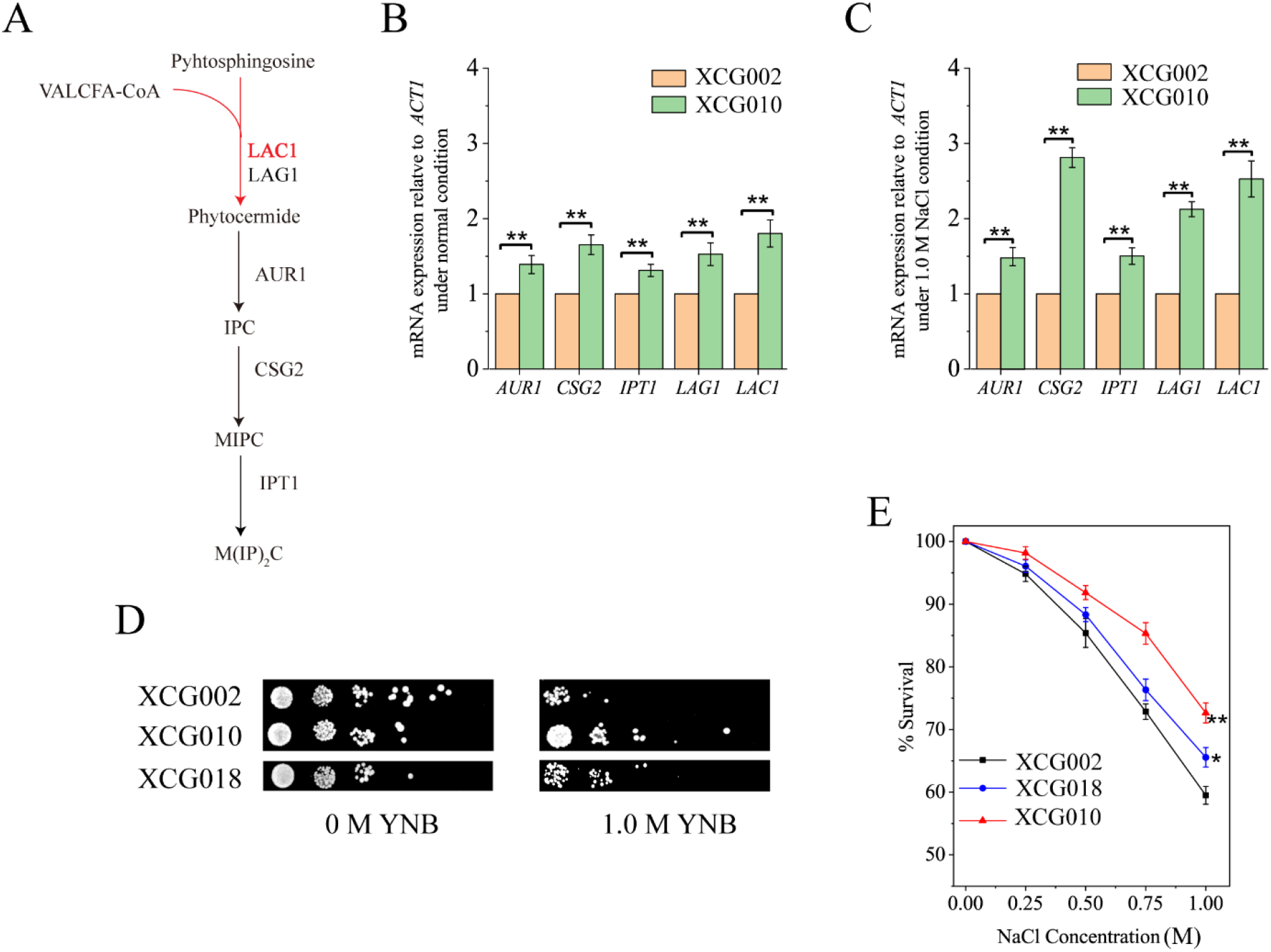
Complex sphingolipids is one key osmotic responsive process to improve osmotic tolerance. (A) Complex sphingolipid biosynthesis pathway. (B) The mRNA level of the complex sphingolipid biosynthesis genes in strain XCG002 and XCG010 under 0 M NaCl. (C) The mRNA level of complex sphingolipid biosynthesis genes in strain XCG002 and XCG010 under 1.0 M NaCl. (D) Strains XCG002, XCG010 and XCG018 were spotted on plates containing with or without 1.0 M NaCl. (E) The survival rates of strains XCG002, XCG010 and XCG018 over of a range of NaCl doses (0.00, 0.25, 0.50, 0.75, 1.00 M). All data are presented as mean values of three independent experiments. Error bars indicate the standard deviations. *, *P* <0.05; **, *P* <0.01; ***, *P* <0.001.

To evaluate whether the inhibition of complex sphingolipid biosynthesis affects the growth of strain XCG010, *LAC1*, which is involved in the synthesis of ceramide from acyl-coenzyme A and phytosphingosine, was deleted to generate strain XCG018. Spot results indicated that, at 1.0 M NaCl, the growth of strain XCG018 was better than that of strain XCG002 but worse than that of strain XCG010 (Fig. 5D). Moreover, cell survival of strain XCG018 (65.6%) was increased by 10.2%, compared with that of strain XCG002 (59.5%) at 1.0 M NaCl (Fig. 5E). These results suggested an increasing complex sphingolipids was crucial for *S. cerevisiae* osmotic tolerance.

### Increased complex sphingolipid content improved membrane integrity

Effect of complex sphingolipids on membrane integrity was investigated (Fig. 6). Strains XCG002, XCG010 and XCG018 were treated with 0 M or 1.0 M NaCl for 4 h and subjected to SYTOX green and FM4-64 uptake analysis. At 0 M NaCl, scanning electron microscope analyzed that almost all cells of strains XCG002, XCG010 and XCG018 exhibited integral membranes (Fig. 6A). At 1.0 M NaCl, some cells of strains XCG002, XCG010 and XCG018 displayed damaged membranes, where XCG010 showed the least, XCG018 showed the second least and XCG002 showed the most number of cells with damaged membranes (Fig. 6B). These cells of strains XCG002, XCG010 and XCG018 were further assayed using flow cytometry. At 0 M NaCl, the percentage of cells with integral membranes in strains XCG010 and XCG018 was similar to that of strain XCG002. While the percentage of cells with integral membranes in strain XCG010 and XCG018 were 85.7% and 76.6%, respectively, which were increases amounting to 24.4% and 11.2%, compared with that of strain XCG002 (68.9%). These results suggested an increasing complex sphingolipid content improved membrane integrity.

**FIG. 6.**
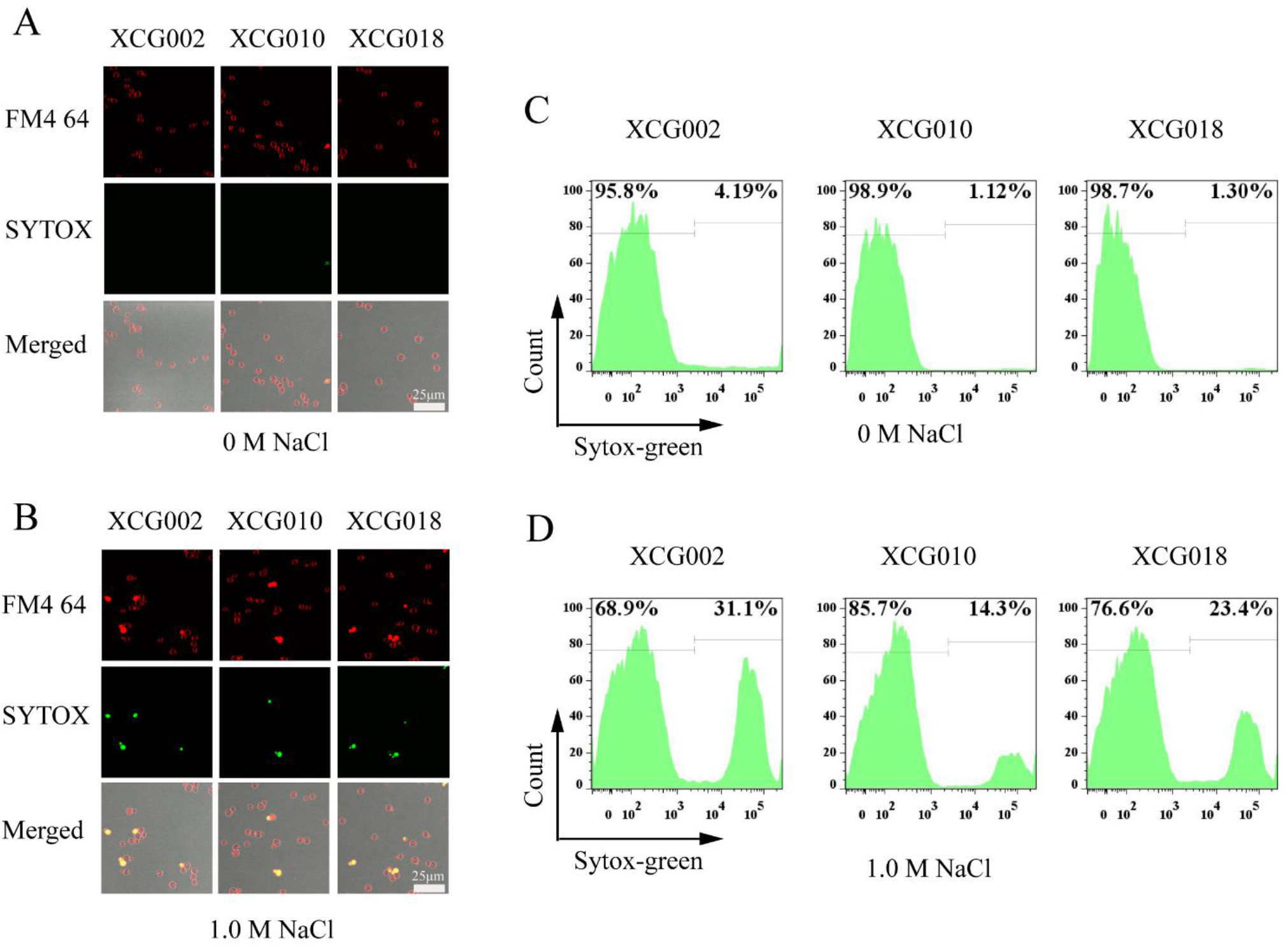
Increased complex sphingolipids changed membrane integrity. (A and B) Scanning electron microscope analysis of membrane integrity in the XCG002, XCG010 and XCG018 cells at 0 M (A) or 1.0 M NaCl. Under the view of a confocal fluorescence microscope, all cells showed red fluorescence with integral membrane while only cells with damaged membrane showed green fluorescence. Cell with damaged membrane can be stained by SYTOX green and cell with integral or damaged membrane all can be stained by FM4-64. Scale is 25 μm. (C and D) Flow cytometry analysis of membrane integrity in strains XCG002, XCG010 and XCG018 cells at 0 Mor 1.0 M NaCl.

## DISCUSSION

In this study, ALE was used to obtain an osmotic tolerant strain, XCG001, where RNA-seq analysis of mutant XCG001 and the wild-type strain was used to identify a key gene, *ELO2*, associated with osmotic tolerance. Furthermore, overexpression of *ELO2* changed the composition of complex sphingolipids and increased the content of complex sphingolipids with longer acyl chain. As a result, membrane integrity was increased and the osmotic resistance was enhanced. This study provides a novel strategy to manipulate membrane complex sphingolipids to increase membrane integrity and osmotic tolerance.

In this study, RNA-seq analysis of mutant XCG001 and the wild-type strain showed that *ELO2* was associated with osmotic tolerance. ALE is a very efficient way to improve industrial strain phenotypes (27, 28). For example, ALE was used to increase the specific growth rate for some genes deletion *S. cerevisiae* or genome-reduced *E.coli* with glucose as energy sources (29, 30), or to improve glycerol assimilation ability of *S. cerevisiae* (31), or enhance *Schizochytrium* sp. tolerance to high salinity stress (32). After securing ALE strains, an important objective was to further identify the targets for genetic modification. Three omics tools were applied for this purpose as follows: (i) Genomics: The growth of pyruvate metabolism modified *S. cerevisiae* was recovered to wild-type strain levels through ALE, and *MED2*^*432Y^ and *GPD1*^W71*^ of 17 mutant genes, which were found in the genome sequence of 7 strains from the evolved *S. cerevisiae*, may play a key role in regulating cell growth, as a result the growth of a *MED2*^*432Y^ and *GPD1*^W71*^ double-mutant strain was similar to the wild-type strain (29); (ii) Transcriptome: The specific growth rate of *S. cerevisiae* on glycerol was increased via ALE, and the transcriptome data revealed that genes, which were related to the tricarboxylic acid cycle and oxidative phosphorylation, contributed to the increased specific growth rate (31). As a result, overexpression of *HAP4*, which is involved in the TCA cycle, and *STL1*, which encodes a glycerol/H^+^ symporter, improved the growth of *S. cerevisiae*; (iii) Metabolomics: An ionic liquids tolerant *Yarrowia lipolytica* strain was obtained through ALE, and metabolomics analysis showed sterols played a key role in ionic liquids tolerance, thus, a sterols transcription factor was overexpressed to enhance ionic liquids tolerance (17).

Overexpression of Elo2, a fatty acid elongase that catalyzes C16-carbon fatty acids to C22 (33), may change lipid composition, including fatty acids, phospholipids, and complex sphingolipids. Sphingolipids play an important role in physiological functions by regulating cell growth and responding to environmental stress (34). Sphingolipids, which are signaling molecules, modulate cellular functions and fate, including cell division, cell death, lifespan, and autophagy (14, 21, 34). Sphingolipid content in the plasma membrane can be manipulated to help cells tolerate stress (22, 24, 35). For example, endogenous sphingolipids were accumulated when mouse sphingomyelin synthase 1 (Sms1) was expressed in yeast, as a result of which tolerance of the strain to oxidation, osmotic, and temperature stresses were improved (22). Furthermore, lipid composition and content may be changed by metabolic pathway genes, harsh environmental conditions, and transcription factors (14, 36). Manipulating lipid biosynthesis genes changed lipid content (37). *ELO3*, an *ELO2* paralog, also regulates sphingolipid composition by elongating of the C18 acyl chain up to C26 (33). Environmental or chemical stresses affect lipid metabolism, which plays a role in maintaining membrane homeostasis and cell growth, a case in point is the membrane unsaturated fatty acids to saturated fatty acids ratio being increased under high-pressure homogenization stress, which enables the strain to avoid damage (36). Transcription factors, such as Mga2 and Upc2 that enable changes in the expression of lipid biosynthesis genes may change lipid composition indirectly (18, 38).

Enhancement of the complex sphingolipids content increases membrane integrity and osmotic tolerance of *S. cerevisiae*. Membrane integrity could be enhanced by engineering membrane components including: (i) Transporter proteins: Membrane integrity could be modified by transporter proteins. For example, when sugar and ion transporter, OmpF, was deleted, and the long chain fatty acids transporter, FadL, was overexpressed in *E. coli*, membrane integrity was enhanced and the fatty acid titer improved (39); (ii) Phospholipids: Membrane integrity can also be altered by modifying the distribution of phospholipid head groups, by adjusting phospholipid saturation, and by altering phospholipid acyl chain length (14, 40). For instance, when pssA, a phosphatidylserine synthase, was overexpressed, PE content increased and membrane integrity was enhanced, as a result of which bio-renewable fuel tolerance and titer was improved (12); (iii) Sterols: Sterol serves as an essential cell membrane component, where deletion of CgMed3AB decreased lanosterol, zymosterol, fecosterol, and ergosterol content, thereby reducing the membrane integrity of *Candida glabrata* (19); (iv) Sphingolipids: Sphingolipid metabolism plays an important role in membrane integrity (41). When sphingolipid biosynthesis genes were deleted in *S. cerevisiae*, the resultant strains exhibited resistance to amphipathic peptidomimetic LTX109, which decreased membrane integrity (41). That study did not demonstrate whether deletion of sphingolipid biosynthesis genes increases membrane integrity directly.

## MATERIALS and METHODS

### Strains and Media

All *S. cerevisiae* strains and plasmids used in this study are listed in Table 1. Plasmids pY131 and pY132 were constructed by replacing the promoter P*_TEF_* of pY13 plasmid with P*_ADH1_* and P*_TDH3_*, respectively. Overexpression strains were constructed using pY13, pY131 and pY132 plasmids carrying the target genes. All plasmids were transformed into yeast cells using the lithium acetate transformation method. Homologous recombination was used for gene *lac1* deletion. The *LEU2* marker, the upstream and downstream regions of the target gene open reading frame were fused by fusion-PCR, and the PCR products were transformed into yeast cells using the lithium acetate transformation method. All primers used in this study are listed in Table 2. Yeast was cultivated in yeast extract peptone dextrose (YPD) medium and yeast nitrogen base (YNB) medium at 30°C with shaking at 200 rpm.

**Table 1.**
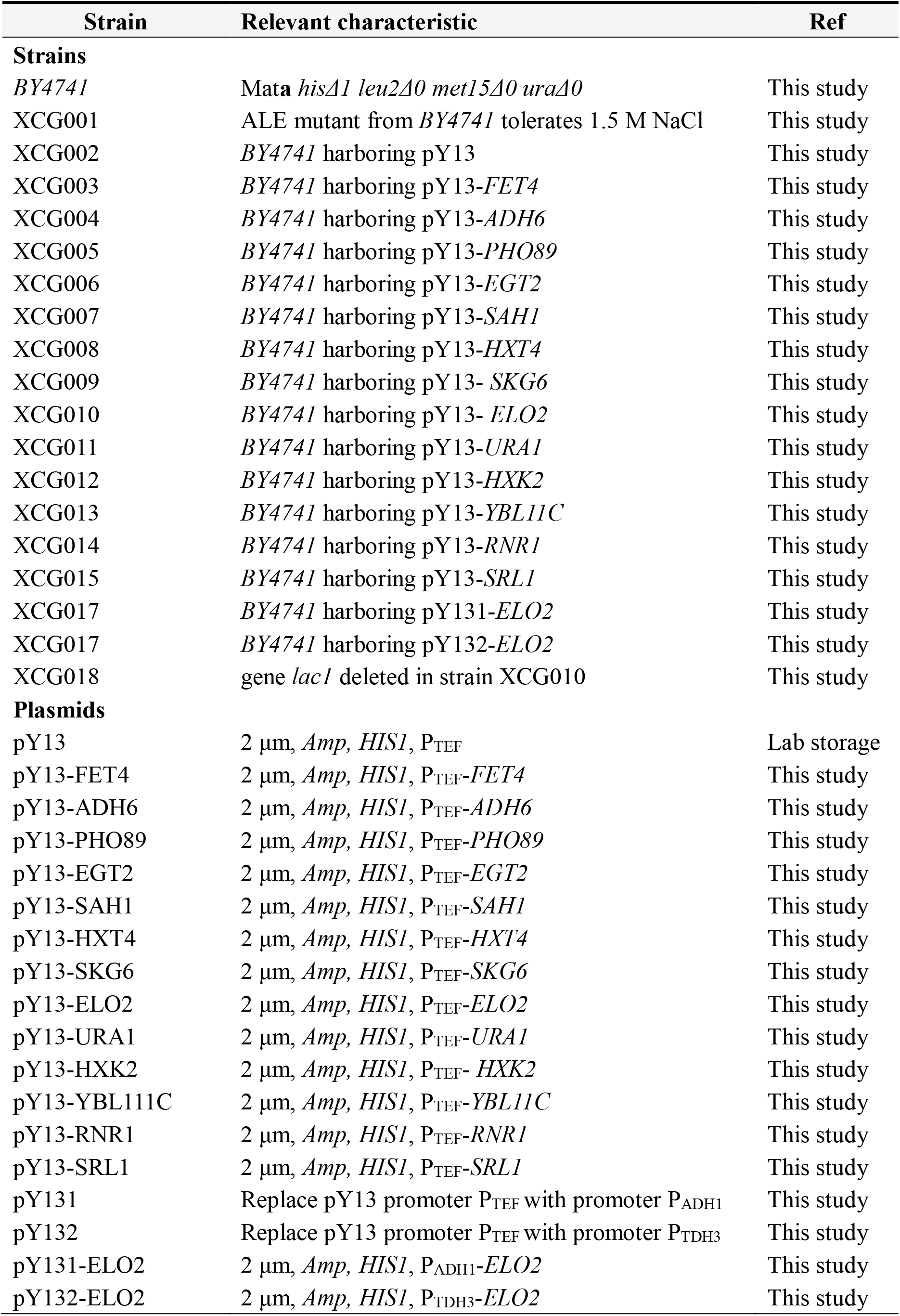
Strains used in this study

**Table 2.**
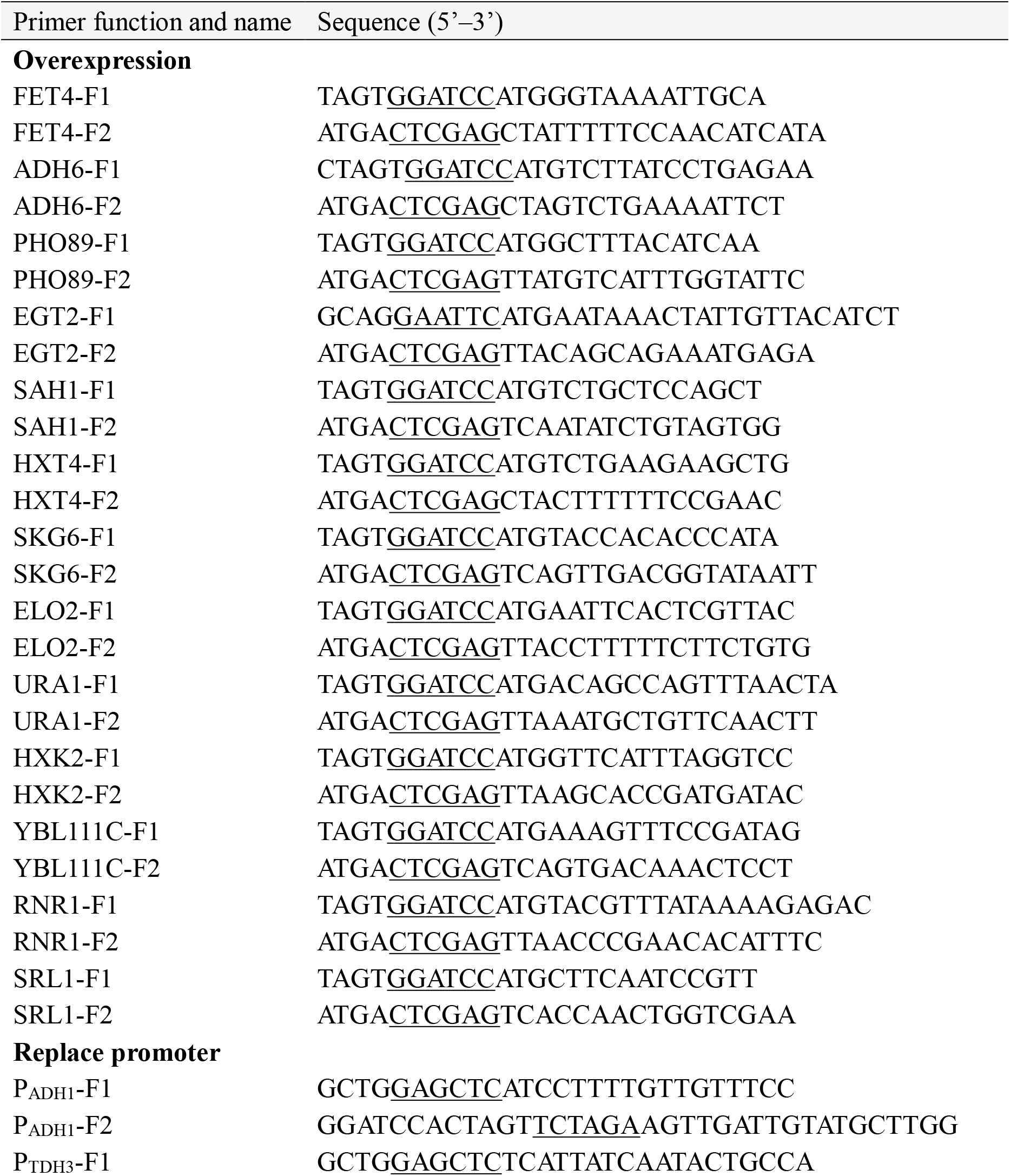
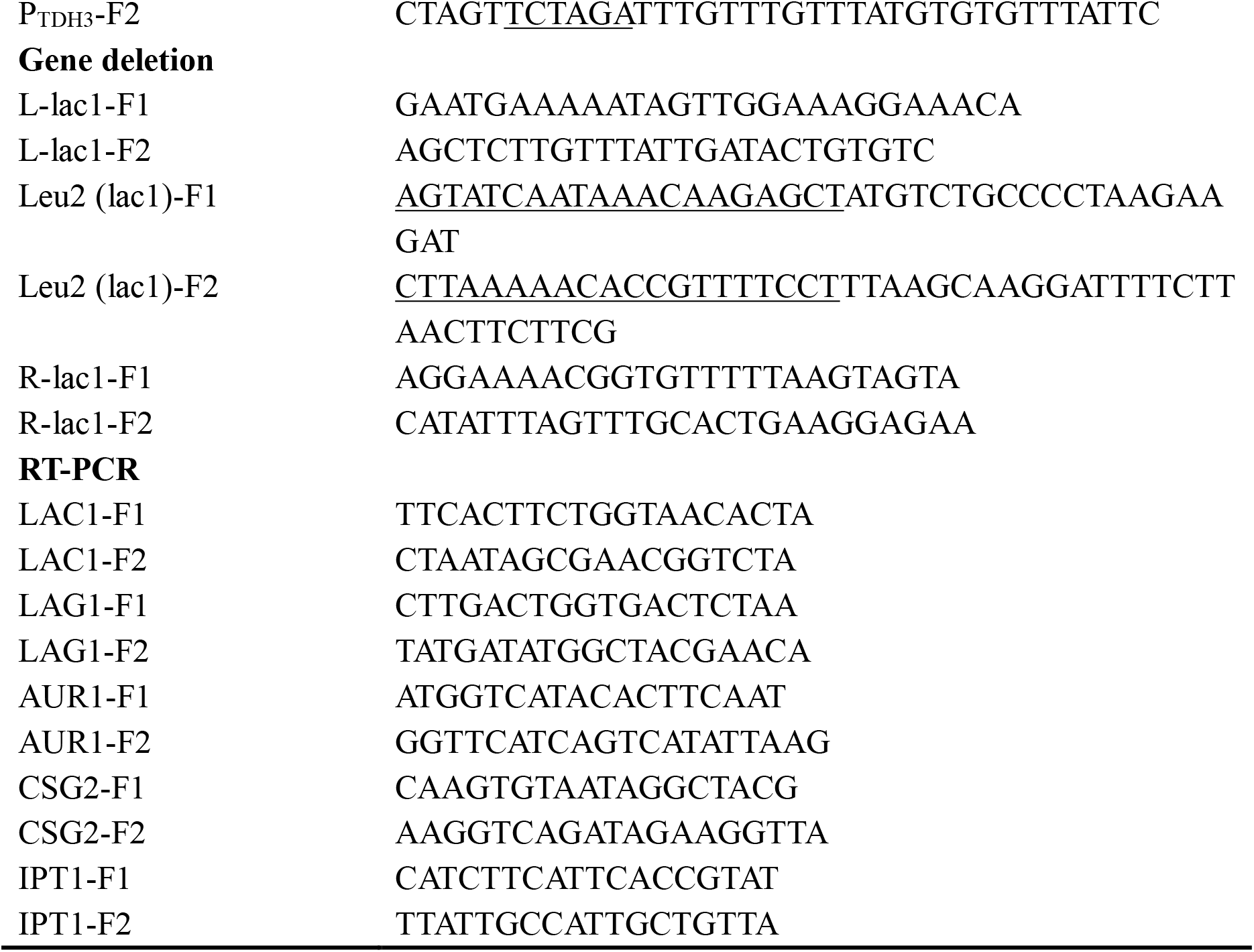
Primer used in this study

### Adaptive laboratory evolution

*S. cerevisiae BY4741* was cultivated in 25 ml of YNB medium with histidine, leucine, methionine uracil and the increasing salt concentrations in a 100-ml flask. When optical density at 600 (OD_600_) reached around 4, the strain was transferred to new salt medium with an initial OD_600_ of approximately 0.1. The concentration of salt was increased when the maximum specific growth rate about 0.3.

### Spot assay

Yeast cells were cultivated in logarithmic (log) phase and diluted to OD_600 nm_ of 1.0. Aliquots (4 μl) of 10-fold serial dilutions were spotted onto YNB agar plates with or without the indicated concentration of NaCl. Growth was assessed after incubation for 2 to 4 days at 30°C.

### IC_50_, Growth curve and Survival

Maximum exponential growth rates of yeast in YNB supplemented with increasing salt concentrations. The half-maximal inhibitory concentration (IC_50_) was calculated by fitting a Hill-type model to the data (solid line). Data points and error bars represent mean and s.d. of three biological replicates. To test growth curve of *S. cerevisiae* at different concentration of NaCl, cells were cultivated in log phase and diluted to a fresh YNB medium with OD_600_ of 0.1 at different concentration of NaCl. The OD_600_ values were recorded though taking curatives at regular time intervals. Cells survival was assessed by log-phase cells were treated with various concentrations of NaCl for 1 h at 30°C with shaking at 200 rpm. Next, cells were diluted and plated on YNB medium plates with various concentrations of NaCl. After incubation for 2 to 4 days at 30°C, the surviving colonies were counted. The survival rates are expressed relative to that of untreated cells of the corresponding strain.

### Transcriptome analysis

The wild type strain and mutant XCG001 were cultured in the log phase at 0 M and 1.5 M NaCl. The collected strains were frozen at −80°C and sent to the Genewiz Institute for RNA extraction and global gene analysis.

### qRT-PCR analysis

Total RNA was extracted using MiniBEST universal RNA extraction kit and 1 μg was taken to synthesize cDNA using the PrimeScript II 1st-strand cDNA synthesis kit (TaKaRa, Japan). The cDNA mixture was diluted to about 100 ng/μl and used as the template for the gene expression level analysis by qRT-PCR. qRT-PCR was performed with TB Green Premix Ex Taq (TaKaRa Bio) using an iQ5 continuous fluorescence detec0tor system (Bio-Rad, Hercules, CA). Data were normalized to that of β-actin gene ACT1. The primer sequences for qRT-PCR are listed in Table 2.

### Fatty acids analysis

Fatty acids of yeast was extracted using NaOH-methanol-distilled water solution (3:10:10, wt/vol/vol) and freeze-dried. Then dried sample was treated with 2 ml boron trifluoride (BF3)- methanol (12:88, vol/vol) and produce fatty acid methyl esters, as described previously (42). Last, samples were analyzed by gas chromatography (GC) with a polyethylene glycol capillary column eluted at a flow rate of 29.6 ml/min and the column pressure of 63.4 kPa (43). Data analysis was based on the following standard Supelco37 (sigam, USA, CAS:47885-U). Fatty acid was ensured by order of Supelco 37 retention time and weight percentage of each fatty acid accounts for Supelco 37 (Fig. S3).

### Phospholipid measure

Phospholipids was extracted from the freeze-dried samples using chloroform-methanol as described previously (44). Dried phospholipids were obtained under a nitrogen stream and reconstituted in chloroform-methanol (1:1, vol/vol). Samples were analyzed by ultrahigh-performance liquid chromatography tandem mass spectrometry (UPLC-MS; Waters, USA) and a CORTECS UPLC hydrophilic interaction liquid chromatography (HILIC) column (2.1 by 150 mm; inner diameter, 1.6-μm) with gradient elution at 45°C and a rate of infusion of 0.3 ml·min^−1^.

### Complex sphingolipid measure

Strains were cultured in YNB medium with or without 1.0 M NaCl for 6-8 h, and washed with PBS. The cell pellets were lysed in PBS by bead-beating mechanical disruption at 4°C. The supernatants were then extracted with chloroform-methanol (2:1, vol/vol) at a final ratio of 20% (v/v). Centrifuge using a refrigerated centrifuge at 4°C to obtain the supernatant. The extracts were evaporated to dryness under nitrogen at room temperature and stored at - 80°C. The dried samples were sent to the Profleader Institute for complex sphingolipids analysis and solubilized in dichloromethane-methanol (2:1,vol/vol) before analysis by UHPLC-QTOF-MS (Agilent) analysis (Fig. S4).

### Cell membrane integrity analysis

Cell membrane integrity was analyzed by microscopy and flow cytometry. For microscopy analysis of cell membrane integrity, the log phase cells were treated with 0 M and 1.0 M NaCl for 4 h and washed with PBS two twice. Then, samples were subjected to SYTOX green and FM4-64 uptake for 20 mins and placed on a microscope slide, covered with a coverslip. Images were acquired using a Nikon ECLIPSE 80i microscope equipped with a Nikon DS-Ri1 camera. For flow cytometry of cell membrane integrity, 10,000 counts of stained cells were recorded using a 0.5 mL s-1 flow rate. All data were exported in FCS3 format and processed using FlowJo software (FlowJo, LLC).

### Statistical analysis

Experimental data are shown as the means ± standard errors of the means (SEM). All quantitative data were analyzed using Student’s t test or one-way analysis of variance (ANOVA). Each experiment was repeated at least three times.

### Accession number(s)

The RNA-seq raw reads were submitted to NCBI under BioProject number PRJNA568205, and the Sequence Read Archive (SRA) entries are SRR10150286, SRR10150285, SRR10150284, SRR10150283.

## ACKNOWLEDGMENTS

This work is supported by the National Key R&D Program of China (2018YFA0901401), the National Natural Science Foundation of China (21808083, 21878126, 21978113), the Key Field R & D Program of Guangdong Province (2019B020218001) and the National First-class Discipline Program of Light Industry Technology and Engineering (LITE2018-08).

G.Z, L.L and J.W designed the research; G.Z and N.Y performed the research; G.Z, Q.L, J.L, X.C and J.W analyzed the research; and G.Z and L.L wrote the paper.

We declare no competing financial interests.

